# O-GlcNAcylation of YTHDF2 antagonizes ERK-dependent phosphorylation and inhibits lung carcinoma

**DOI:** 10.1101/2023.09.09.557012

**Authors:** Jie Li, Wen Zhou, Jianzhi Zhang, Li Ma, Zhuan Lv, Yiqun Geng, Xing Chen, Jing Li

**Affiliations:** Beijing Key Laboratory of DNA Damage Response and College of Life Sciences, Capital Normal University, Beijing 100048, China; College of Chemistry and Molecular Engineering, Beijing National Laboratory for Molecular Sciences, Peking-Tsinghua Center for Life Sciences, Synthetic and Functional Biomolecules Center, and Key Laboratory of Bioorganic Chemistry and Molecular Engineering of Ministry of Education, Peking University, Beijing 100871, China; Key Laboratory of Carcinogenesis and Translational Research, Ministry of Education, Department of Thoracic Surgery II, Peking University Cancer Hospital & Institute, Beijing 100052, China; State Key Laboratory of Bioactive Substance and Function of Natural Medicines, Institute of Materia Medica, Chinese Academy of Medical Sciences and Peking Union Medical College, Beijing 100050, China

**Keywords:** O-GlcNAc, m^6^A, c-Myc, ERK, lung carcinoma

## Abstract

The intracellular O-linked N-acetylglucosamine (O-GlcNAc) glycosylation mediates many signal transduction events and regulates tumorigenesis. Previously the RNA N6-methyladenosine (m^6^A) reader, YTH (YT521-B homology) domain 2 (YTHDF2), has been shown to be O-GlcNAcylated on Ser-263 during Hepatitis B virus (HBV) infection and promote HBV-related hepatocellular carcinoma. Herein we mapped YTHDF2 O-GlcNAcylation at Thr-49 via electron-transfer dissociation mass spectrometry under unperturbed conditions. We show that YTHDF2 Thr-49 O-GlcNAcylation antagonizes Extracellular-signal regulated kinase (ERK)-dependent phosphorylation at Ser-39 and promotes YTHDF2 degradation. The downstream signaling pathway of YTHDF2 in lung carcinoma are thus upregulated, which leads to the downregulation of c-Myc. We further used mouse xenograft models to show that YTHDF2-T49A mutants increased lung cancer mass and size. Our work reveals a key role of YTHDF2 O-GlcNAcylation in tumorigenesis and suggests that O-GlcNAcylation exerts distinct functions under different biological stress.

## INTRODUCTION

O-linked N-acetylglucosamine (O-GlcNAc) glycosylation is a post-translational modification (PTM) that involves decorating the Ser/Thr residues with the O-GlcNAc moiety (1,2). It is catalyzed by the O-GlcNAc transferase (OGT) and removed by the O-GlcNAcase (OGA). Decades of glycobiology research have witnessed emerging chemical and biological tools (3,4) that have demonstrated that O-GlcNAc occurs on about 5 000 proteins and regulates tumorigenesis and immunotherapy (5), ferroptosis (6), protein aggregation and phase separation (7), stem cell biology (8), liver metabolism (9), neurodevelopment and various stress response pathways (10).

RNA N6-methyladenosine (m^6^A) is one of the ∼140 types modifications in RNA, especially in mRNAs and lncRNAs (11). It is the most abundant internal modifications that have lots of therapeutic potential (12,13). The regulation of m^6^A is intricate, which includes its writers, readers and erasers (14). Among the readers, the YTH (YT521-B homology) family proteins have caught a lot of attention (15), because not only they all contain the conserved YTH domain that binds m^6^A, but also they are subject to many PTMs that finetune their biological functions under physiology and pathology conditions.

Our lab has been intensively studying the relationship between O-GlcNAc and m^6^A readers, including YT521-B homology (YTH) domain 1 (YTHDF1) and YTH domain-containing protein 1 (YTHDC1). We have shown that O-GlcNAcylation of YTHDF1 promotes its cytosolic localization and enhances downstream target expression (16). We also demonstrated that DNA damage induces O-GlcNAcylation on YTHDC1, which promotes its binding with m^6^A and subsequent homologous recombination (17). In this work we focused on YTH domain 2 (YTHDF2) and aimed to elucidate the role of O-GlcNAc in regulating YTHDF2.

YTHDF2 is a reader that regulates m^6^A mRNA decay (18). Among its various roles in human diseases, it has a particular role in tumor (19,20). In acute myeloid leukemia, YTHDF2 is often overexpressed and decreases the stability of many m^6^A transcripts, whose function is required for leukemic stem cell function (21). In multiple myeloma, YTHDF2 overexpression correlates with poor prognosis, and the underlying mechanism is that YTHDF2 inhibits MAP2K2/p-ERK (22). In lung adenocarcinoma, YTHDF2 positively regulates tumorigenesis by promoting the mRNA decay of AXIN1, which negatively regulates the Wnt/β-catenin pathway (23). Recently YTHDF2 inhibition has also been shown to improve radiotherapy efficacy (24).

Due to its pivotal role in tumorigenesis, YTHDF2 is subject to many PTMs. YTHDF2 protein stability is dependent on cyclin dependent kinase (CDK1), although a direct phosphorylation reaction is yet to be revealed (25). YTHDF2 is phosphorylated by the EGFR/SRC/ERK signaling pathway at Ser-39 and Thr-381 and thus stabilized and promotes invasive glioblastoma (26). Hypoxia induces small ubiquitin-like modifier (SUMO) modification of YTHDF2 at Lys-571, which increases its binding with m^6^A and results in cancer (27). Upon hepatitis B virus (HBV) infection, YTHDF2 is O-GlcNAcylated at Ser-263 to enhance its stability, and downstream target expression (including minichromosome maintenance protein 2 and 5) and HBV-related hepatocellular carcinoma (28). YTHDF2 is ubiquitination by STIP1 homology and U-box-containing protein 1 (STUB1) to inhibit Sorafenib resistance in hepatocellular carcinoma (HCC) (29).

In this paper, we found that YTHDF2 is O-GlcNAcylated during unperturbed conditions. Using electron-transfer dissociation (ETD) mass spectrometry (MS), we identified Thr-49 as the O-GlcNAc site. Thr-49 O-GlcNAcylation antagonizes ERK-dependent phosphorylation at Ser-39, probably due to its close proximity. We show that YTHDF2 O-GlcNAcylation decreases its abundance. In lung carcinoma cells YTHDF2 O-GlcNAcylation downregulates c-Myc, possibly through the AXIN1/Wnt/β-catenin pathway, thus inhibits tumor progression in mouse xenograft models. Our work reveals that O-GlcNAcylation could occur on distinct sites in different biological settings, and YTHDF2 O-GlcNAcylation provides new therapeutic potential for lung carcinoma.

## Results

### YTHDF2 is O-GlcNAcylated at Thr-49 under unperturbed conditions

To examine if YTHDF2 is O-GlcNAcylated under basal conditions, we first tested the interaction between YTHDF2 and OGT. Cell lysates were subject to immunoprecipitation with anti-OGT antibodies, and then immunoblotted (IBed) with anti-YTHDF2 and anti-OGT antibodies (Fig. 1A). The results showed that endogenous YTHDF2 co-immunoprecipitates (coIPs) with OGT. Then cells were transfected with Myc-OGT and HA-YTHDF2 plasmids to examine interaction between exogenously expressed proteins. Again HA-YTHDF2 coIPs with Myc-OGT (Fig.1B). Then GST-pulldown experiments were carried out. Cells were transfected with HA-YTHDF2, and the cellular lysates were incubated with recombinant GST-OGT proteins. As shown in Fig. 1C, GST-OGT pulled-down HA-YTHDF2, suggesting the physical association between OGT and YTHDF2 without viral invasion.

**Fig. 1.**
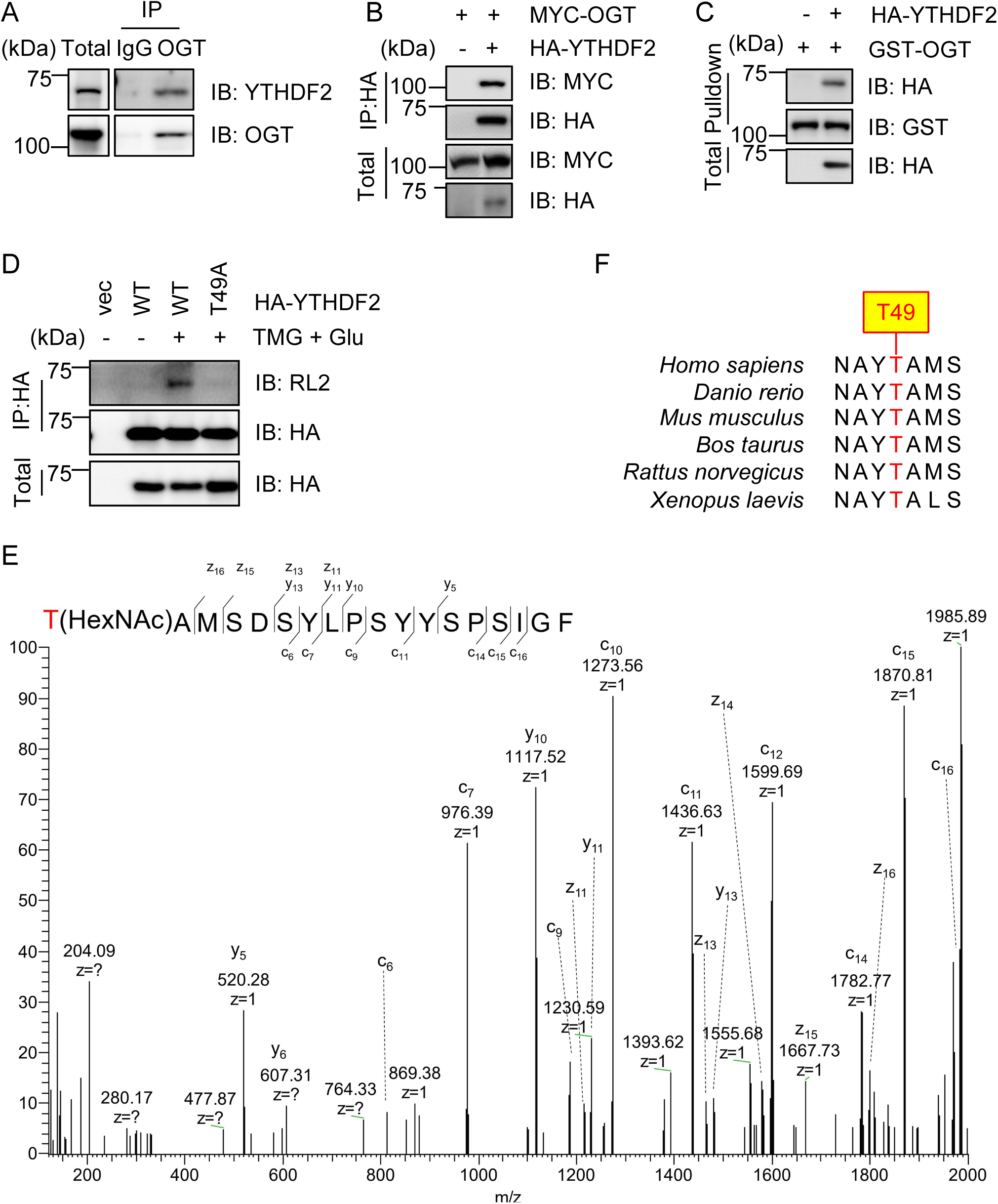
YTHDF2 is O-GlcNAcylated at Thr-49. A, 293T cell lysates were immunoprecipitated with anti-OGT antibodies and immunoblotted with anti-YTHDF2 and anti-OGT antibodies. B, Cells were transfected with HA-YTHDF2 and Myc-OGT plasmids. The cell lysates were subject to immunoprecipitation and immunoblotting with the antibodies indicated. C, Cells were transfected with HA-YTHDF2 or vector controls and the cellular lysates were incubated with recombinant GST-OGT proteins purified from *E. coli.* D, Cells were transfected with HA-YTHDF2-WT or -T49A plasmids, treated or untreated with the OGA inhibitor Thiamet-G (TMG) and glucose to enrich for O-GlcNAcylation. Then the cell lysates were immunoprecipitated with anti-HA antibodies and immunoblotted with anti-O-GlcNAc RL2 antibodies. E, Electron Transfer Dissociation (ETD) mass spectrometry identified that Thr-49 is O-GlcNAcylated. F, Thr-49 is evolutionarily conserved among different species. All Western blots were repeated for at least three times.

We then assessed YTHDF2 O-GlcNAcylation. Cells were treated with Thiamet-G (TMG, OGA inhibitor) plus glucose to enrich for O-GlcNAcylation, as previously described (30). As shown in Fig. 1D, YTHDF2 shows robust O-GlcNAc signaling, indicated by RL2 (O-GlcNAc antibody) staining. The immunoprecipitates were then subject to electron-transfer dissociation (ETD) mass spectrometry (MS) analysis, and a single O-GlcNAc site (Thr-49) was revealed (Fig. 1E), which is quite distinct from the previously identified Ser-263 under HBV infection. We generated T49A mutants accordingly, and the mutant almost abolished O-GlcNAc signals (Fig. 1D). Thr-49 is also conserved among different species (Fig. 1F). These results suggest that YTHDF2 is O-GlcNAcylated under basal conditions.

### YTHDF2 O-GlcNAcylation antagonizes ERK-dependent phosphorylation

As YTHDF2 has been shown to be phosphorylated by ERK at Ser-39 (26), which is quite close to Thr-49, we wonder whether there is a crosstalk between these two PTMs. Thus, we manufactured a phospho-specific antibody targeting p-S39. We first tested the specificity and efficiency of the phospho-antibody (Fig. 2A-B). Cells were transfected with YTHDF2-wild type (WT) or -S39A plasmids, and the p-S39 antibody did not show any signal in the YTHDF2-S39A extracts (Fig. 2A). Then YTHDF2-WT lysates were treated with phosphatase, which again abolished the p-S39 signal (Fig. 2B), suggesting that the antibody is specific.

**Fig. 2.**
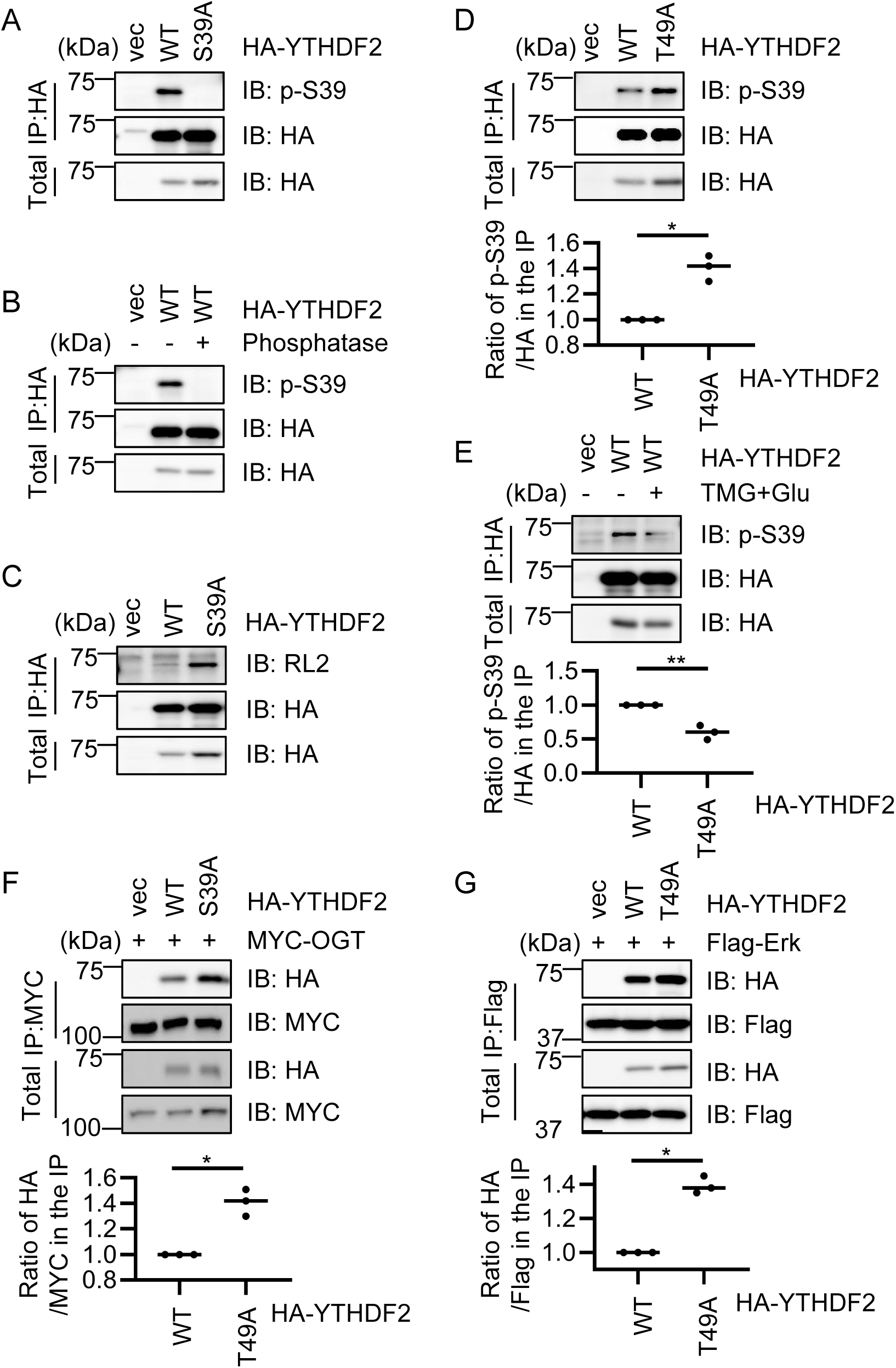
O-GlcNAcylation of YTHDF2 at Thr-49 antagonizes ERK-dependent phosphorylation at Ser-39. A-B, A rabbit anti-pS39 antibody was generated and its specificity was detected. 293T cells were transfected with HA-YTHDF2-WT or HA-YTHDF2-S39A, and the lysates were immunoprecipitated with anti-HA antibodies and immunoblotted with anti-pS39 antibodies (A). Cells were transfected with HA-YTHDF2 plasmids. The anti-HA immunoprecipitates were subject to λ phosphatase (PPase) treatment or left untreated. They were then subjected to immunoblotting with anti-pS39 antibodies (B). C-D, Cells were transfected with HA-YTHDF2-WT or HA-YTHDF2-S39A (C), or -T49A (D) plasmids, and then the lysates were immunoprecipitated with anti-HA antibodies and immunoblotted with indicated antibodies. E, Cells were transfected with HA-YTHDF2-WT plasmids, treated or untreated with the OGA inhibitor Thiamet-G (TMG) and glucose to enrich for O-GlcNAcylation. Then the cell lysates were immunoprecipitated with anti-HA antibodies and immunoblotted with anti-pS39 antibodies. F-G, Cells were transfected with Myc-OGT together with HA-YTHDF2-WT or -S39A (F), or Flag-Erk together with HA-YTHDF2-WT and -T49A (G), and then the lysates were immunoprecipitated and immunoblotted with the antibodies indicated. Statistics analysis was carried out with Student’s *t*-test. * indicates P < 0.05; ** indicates P < 0.01. All Western blots were repeated for at least three times.

We then assessed the potential crosstalk between Thr-49 O-GlcNAcylation and Ser-39 phosphorylation. Cells were transfected with YTHDF2-WT or -S39A, and the S39A mutant shows a drastic increase of O-GlcNAc signals (Fig. 2C). Cells were also transfected with YTHDF2-WT or T49A, and the p-S39 antibody was used for IB. The T49A mutant increased the p-S39 signals markedly (Fig. 2D). On the contrary, the p-S39 signals were decreased when the cells were treated with TMG plus glucose (Fig. 2E). We further investigated whether the changes in modification levels were due to different binding affinity with the catalyzing enzymes. HA-YTHDF2-S39A increased binding with Myc-OGT (Fig. 2F) and HA-YTHDF2-T49A increased binding with ERK (Fig. 2G). These results suggest that O-GlcNAcylation and phosphorylation counteract against each other.

### O-GlcNAcylation decreases YTHDF2 stability

As YTHDF2 Ser-39 phosphorylation plays a role in stabilizing YTHDF2 (26), we wondered whether Thr-49 O-GlcNAcylation would destabilize YTHDF2. Thus, we tested the ubiquitination levels of YTHDF2-WT and -T49A. As shown in Fig. 3A-B, T49A reduced the ubiquitination levels of YTHDF2. To exclude the possibility that this is caused by mutation itself, we used the OGA plasmid to suppress cellular O-GlcNAcylation (Fig. 3C-D). Consistently, OGA overexpression decreased YTHDF2 ubiquitination levels. Cycloheximide (CHX) pulse-chase experiments were performed, and T49A again increased YTHDF2 stability (Fig. 3E-F). Taken together, O-GlcNAcylation at T49 destabilizes YTHDF2.

**Fig. 3.**
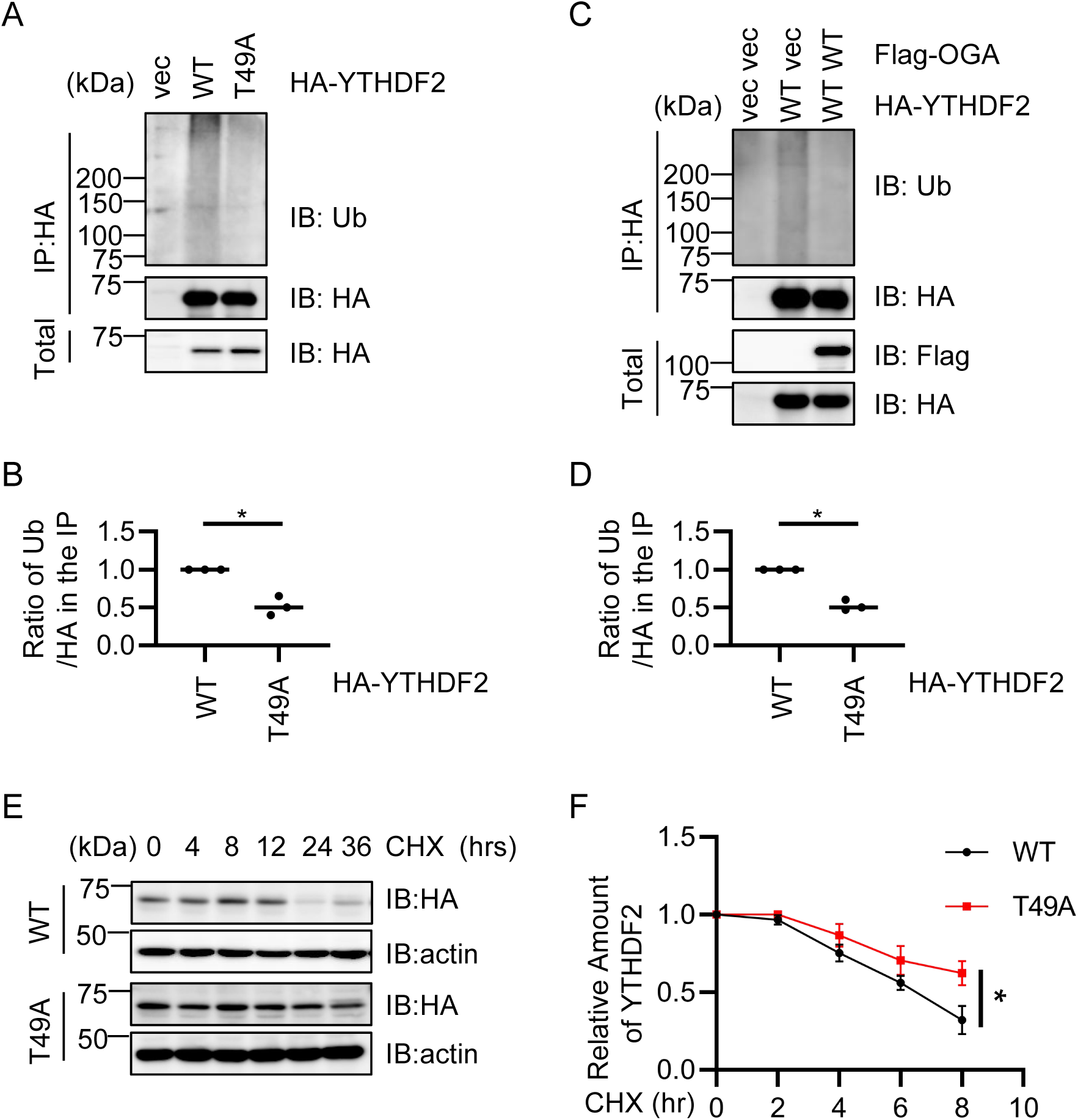
YTHDF2 O-GlcNAcylation promotes ubiquitination. A, 293T cells were transfected with HA-YTHDF2-WT or -T49A plasmids and the lysates were immunoprecipitated with anti-HA antibodies and immunoblotted with anti-Ubiquitin antibodies. B, Quantitation of (A). C, 293T cells were transfected with HA-YTHDF2-WT together with vector or Flag-OGA plasmids and the lysates were immunoprecipitated with anti-HA antibodies and immunoblotted with anti-Ubiquitin antibodies. D, Quantitation of (C). E-F, cycloheximide (CHX) pulse-chase assays. HEK-293T cells were transfected with HA-YTHDF2-WT or -T49A plasmids, then treated with CHX for different durations (E). The quantitation is in (F). Student’s *t*-test was used for statistical analysis in B and D, two-way Anova was used for F. * indicates P < 0.05. All Western blots were repeated for at least three times.

### YTHDF2 O-GlcNAcylation decreases c-Myc via the **β**-catenin pathway in lung carcinoma cells

As YTHDF2 has been demonstrated to exert its function in lung adenocarcinoma through the AXIN/Wnt/β-catenin pathway (23), we examined whether Thr-49 O-GlcNAcylation plays a role. We used both A549 and H1299 lung adenocarcinoma cells, and found that overexpression of T49A markedly increased c-Myc abundance, concomitantly with increased β-catenin levels (Fig. 4 A-D), suggesting that Thr-49 O-GlcNAcylation promotes c-Myc in lung carcinoma.

**Fig. 4.**
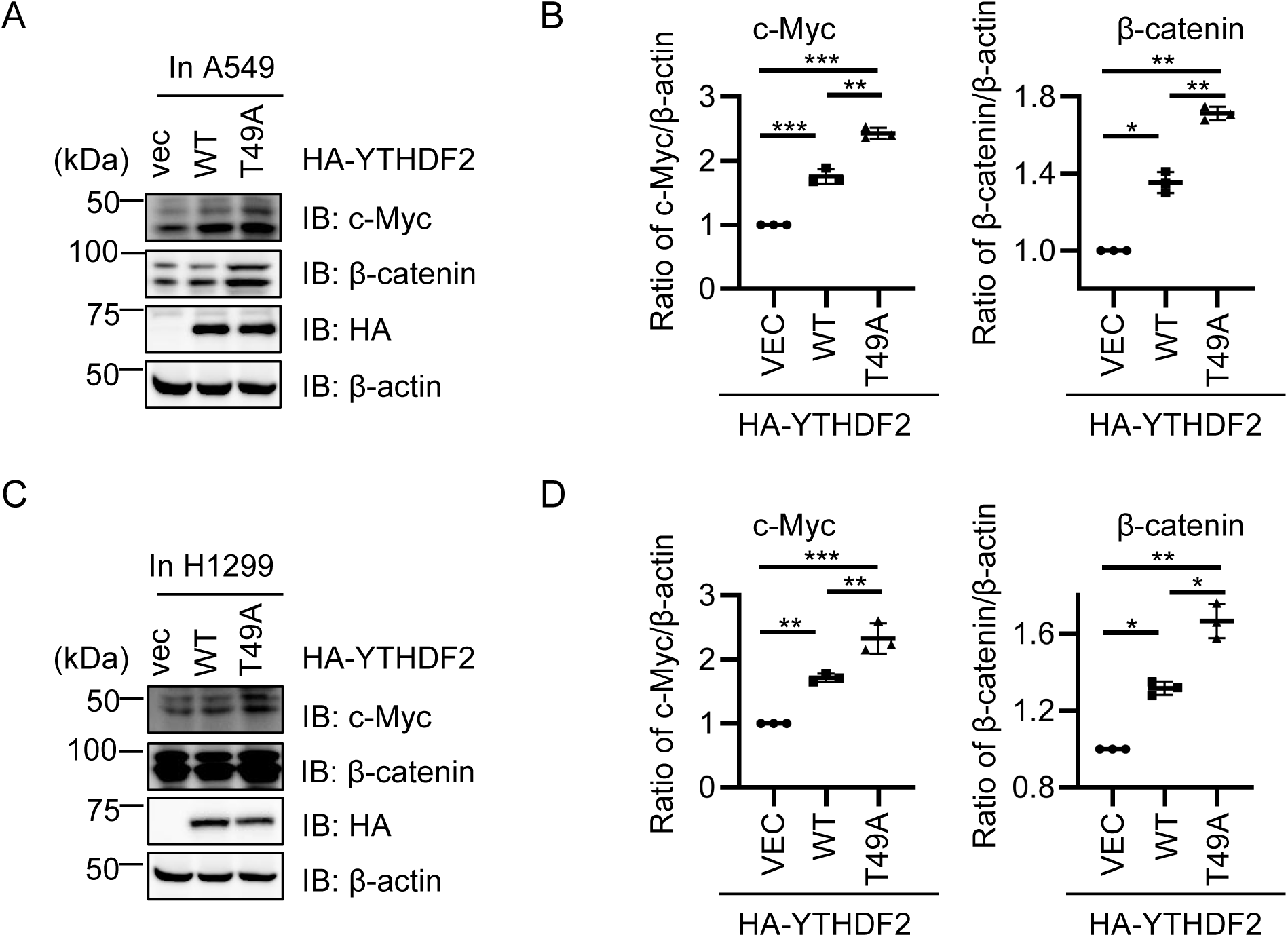
O-GlcNAcylation of YTHDF2 downregulates c-Myc in H1299 and A549 lung cancer cells. A, A549 cells were transfected with HA-YTHDF2-WT or -T49A plasmids and the lysates were immunoblotted with anti-β-catenin and anti-c-Myc antibodies. B, Quantitation of (A). C, A549 cells were transfected with HA-YTHDF2-WT or -T49A plasmids and the lysates were immunoblotted with anti-β-catenin and anti-c-Myc antibodies. D, Quantitation of (C). Statistics analysis were carried out with one-way Anova. * indicates P < 0.05, ** indicates P < 0.01, *** indicates P < 0.001. All Western blots were repeated for at least three times.

### YTHDF2 O-GlcNAcylation downregulates invasion and migration of lung carcinoma cells

To test the effect of YTHDF2 O-GlcNAcylation in cells, we constructed stable transfectants of YTHDF2-WT and -T49A in both A549 and H1299 cells (Fig. 5A). Colony formation assays were carried out and T49A enhanced cellular growth and proliferation (Fig. 5B-C). Trypan blue live cell count assays were performed, and again T49A increased cell growth and proliferation (Fig. 5D-E). We also measured the metastatic capacity by wound healing (Fig. 5F-G) and Transwell invasion (Fig. 5H-I) assays. T49A upregulated cell migration capabilities, suggesting that Thr-49 O-GlcNAcylation attenuates lung carcinoma metastasis.

**Fig. 5.**
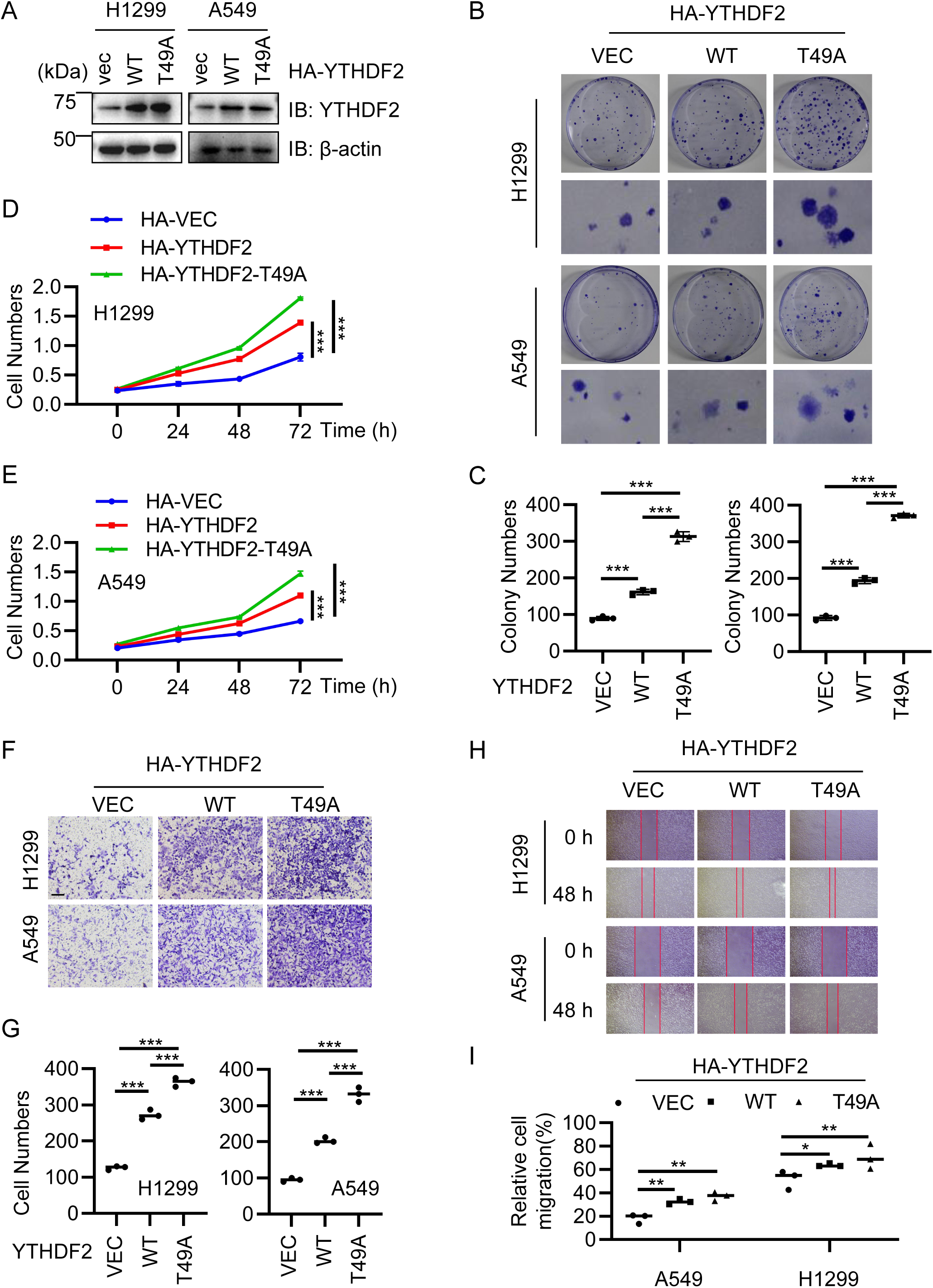
O-GlcNAcylation of YTHDF2 regulates the metastatic capacity of H1299 and A549 lung cancer cells. A, Stable cell lines were generated expressing HA-YTHDF2-WT or -T49A in H1299 and A549 cells. B, Colony formation assays of cells in (A). C, Quantitation of (B). D-E, Cell growth assays of H1299 (D) and A549 (E) cells described in (A). F, Transwell assays of H1299 and A549 cells described in (A). G, Quantitation of (F). H, Wound healing assay of H1299 and A549 cells described in (A). I, Quantitation of (H). Statistics analysis were carried out with two-way Anova for D-E, and one-way Anova for C, G and I. * indicates P < 0.05, ** indicates P < 0.01, and *** indicates P < 0.001.

### YTHDF2 O-GlcNAcylation downregulates lung carcinoma in xenograft mouse models

We then used mouse xenograft models to study Thr-49 O-GlcNAcylation *in vivo*. Stable transfectants were injected into nude mice. After sacrificing the animals, tumors were isolated (Fig. 6A-C), and T49A increased tumor volume and weight than YTHDF2-WT. Taken together, these results suggest a model where O-GlcNAcylation at Thr-49 destabilizes YTHDF2, suppresses lung cancer progression by downregulating c-Myc (Fig. 6D).

**Fig. 6.**
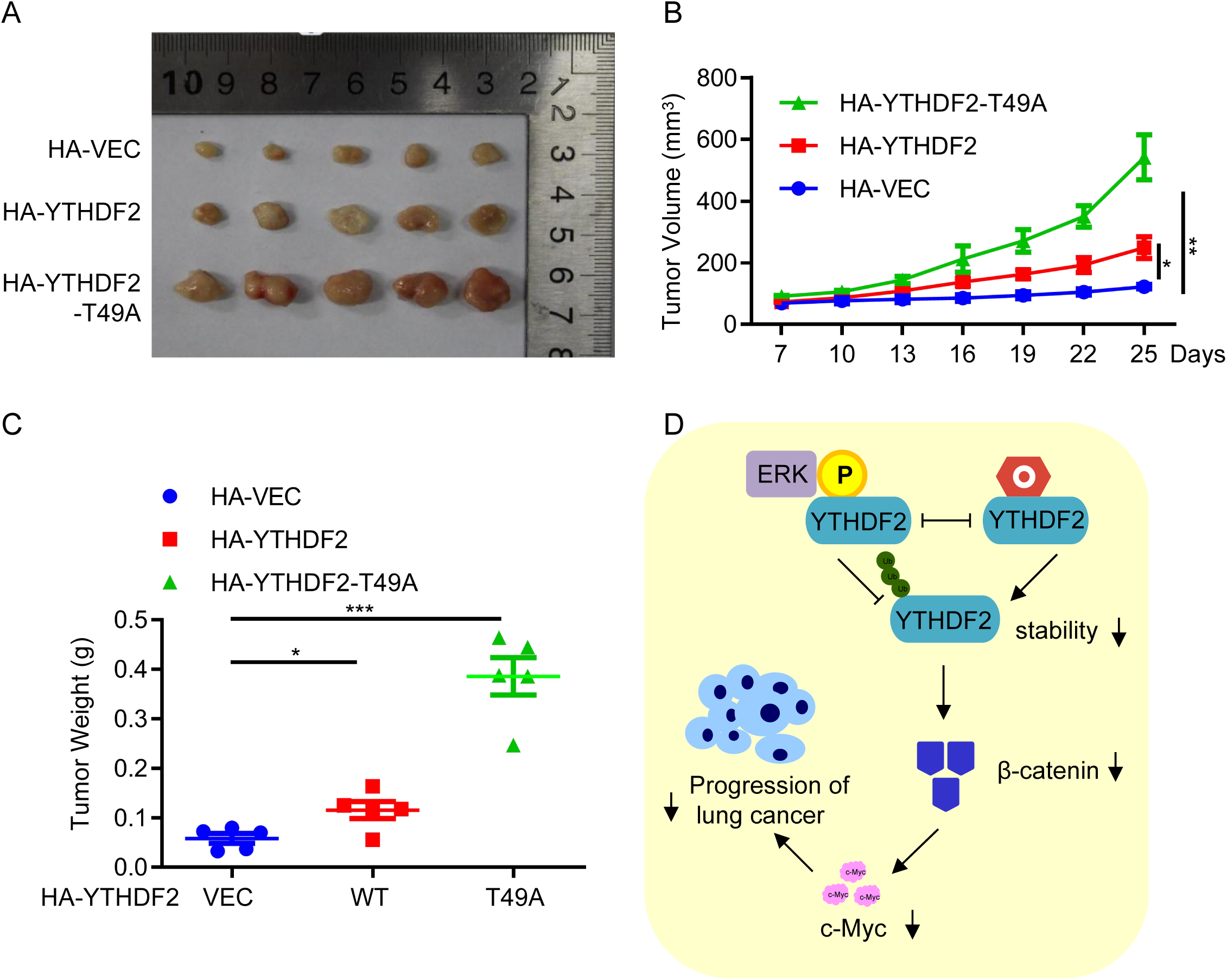
YTHDF2 O-GlcNAcylation inhibits lung cancer. A-C, xenografts in nude mice. HA-YTHDF2-WT and -T49A cells were injected into nude mice. Tumors were photographed after 25 days. Tumor images are in (A), tumor volumes are quantitated in (B) and tumor weights are quantitated in (C). Statistics analysis were carried out with two-way Anova for B, and one-way Anova for C. * indicates P < 0.05, ** indicates P < 0.01, *** indicates P < 0.001. D, Model depicting the role O-GlcNAcylation of YTHDF2: it antagonizes ERK-dependent phosphorylation and inhibits lung arcinoma.

## DISCUSSION

In this work we found that YTHDF2 is O-GlcNAcylated at Thr-49 under basal conditions. Thr-49 O-GlcNAcylation antagonizes Ser-39 phosphorylation by ERK and promotes YTHDF2 ubiquitination and proteasome-mediated degradation. By downregulating c-Myc expression, YTHDF2 O-GlcNAcylation inhibits lung adenocarcinoma (Fig. 6D).

Our work is distinct from previous reports, where O-GlcNAcylation is found to stabilize YTHDF2 upon HBV infection and promote HBV-related hepatocellular carcinoma (28). In our experience, O-GlcNAcylation usually carries out different functions under basal conditions versus viral invasion. For example, under unperturbed conditions UNC51-like kinase-1 (ULK1) is O-GlcNAcylated at Thr-635/Thr-754, which facilitates AMP-activated protein kinase (AMPK)-mediated ULK1 phosphorylation at Ser-555 and Ser-638, and promotes autophagy (31). But upon Human papillomavirus (HPV) infection, ULK1 is O-GlcNAcylated at Ser-409/Ser-410, which antagonizes PKC_α_-mediated phosphorylation at Ser423 and stabilizes ULK1 by counteracting chaperone-mediated autophagy (32). As a nutrient sensor that links stress response with metabolism, perhaps O-GlcNAc responds to virion particles by changing the modified sites and subsequent biological pathways.

Of the three m^6^A readers we investigated (YTHDF1, YTHDC1 and YTHDF2), O-GlcNAc plays distinct roles. YTHDF1 is a nuclear-cytoplasmic shuttling protein, whose nuclear import is inhibited by O-GlcNAcylation. By promoting YTHDF1 downstream targets, such as c-Myc, O-GlcNAc promotes colorectal cancer progression (16). YTHDC1 O-GlcNAcylation is induced by DNA damage. Occurring right on the YTH domain, O-GlcNAc promotes YTHDC1-m^6^A binding and subsequence DNA damage repair by homologous recombination (17). Here we show that YTHDF2 is also O-GlcNAcylated. By promoting YTHDF2 degradation, O-GlcNAc inhibits lung carcinoma. As small noncoding RNAs are recently shown to be modified by sialylated glycans (33), more tools are being developed to enable the discovery of glycoRNAs (34). We think that glycosylation is bound to be found in many RNA metabolism pathways.

### EXPERIMENTAL PROCEDURES

#### Cell culture, antibodies

293T, H1299 and A549 cells were purchased from ATCC. Antibodies were as follows: anti-Myc (PTM Bio, #PTM-5390), anti-c-Myc (ProteinTech, #10828-1-AP), anti-YTHDF2 (proteinTech, #24744-1-AP), anti-Flag (Sigma, #F1084), RL2 (Abcam, #AB2739), anti-GST (Gene Script, #A00865), anti-HA (Bethyl, #A190-108A), anti-OGT (Santa Cruz Biotechnology, #sc-74546), anti-β-actin (Sigma, #A5441), anti-Ubiquitin (PTM Bio, #PTM-1106RM). Anti-YTHDF2-pS39 antibodies were generated using the sequence PYLpSPQAR by Dia-An Biotech, Inc.

#### Immunoprecipitation (IP) and immunoblotting assays

Immunoprecipitation and immunoblotting experiments were performed as described before (35). The following primary antibodies were used for immunoblot: anti-HA (1:5000), anti-FLAG (1:4000), anti-OGT (1:1000), anti-Myc (1:3000), anti-c-Myc (1:1000), anti-YTHDF2 (1:1000), anti-GST (1:3000), RL2 (1:1000), anti-β-actin (1:3000), anti-Ubiquitin (1:1000), anti-pS39 (1:1000). Peroxidase-conjugated secondary antibodies were from JacksonImmuno Research. The ECL detection system (Amersham) was used for immunoblotting. LAS-4000 was employed to detect signals, and quantitated using Multi Gauge software (Fujifilm). All western blots were repeated at least three times.

#### Ubiquitination assays

HEK-293T cells were transfected with plasmids and treated with the proteasome inhibitor MG132 (10 μM) for 12 h. Transfected YTHDF2, and its binding protein were pulled down using the Co-IP assay, and ubiquitination levels were detected using anti-ubiquitin antibodies.

#### IP-phosphatase assay

HEK-293T cells were transfected with HA-YTHDF2 plasmids. The anti-HA immunoprecipitates were subject to λ phosphatase (PPase) treatment or left untreated. Incubation in a water bath at 30°C for 1 h, and then immunoblotted with the antibodies indicated. Lambda PPase was added according to the kit procedure (Lambda PP. NEB #P0753L).

#### Mass Spectrometry

##### Sample Preparation

The gel band pieces were dehydrated in acetonitrile, incubated in 10 mM DTT in 50 mM ammonium bicarbonate at 56 °C for 40 min, incubated in 55 mM iodoacetamide in 50 mM ammonium bicarbonate at ambient temperature for 1 hr in the dark, and dehydrated again. Then the gel pieces were digested in-gel with 2 ng/μL sequencing grade chymotrypsin in 50 mM ammonium bicarbonate overnight at 37 °C. The resulting peptides were extracted twice with 5% formic acid/50% acetonitrile, then vacuum-centrifuged to dryness. All samples were resuspended in 0.1% FA in water prior to LC-MS/MS analysis.

##### LC-MS/MS Parameters

Peptides were separated using a loading column (100 µm × 2 cm) and a C18 separating capillary column (75 µm × 15 cm) packed in-house with Luna 3 μm C18(2) bulk packing material (Phenomenex, USA). The mobile phases (A: water with 0.1% formic acid and B: 80% acetonitrile with 0.1% formic acid) were driven and controlled by a Dionex Ultimate 3000 RPLC nano system (Thermo Fisher Scientific). The LC gradient was held at 2% for the first 8 minutes of the analysis, followed by an increase from 2% to 44% B in 60 minutes, and an increase from 44% to 99% B in 5 minutes.

For the samples analyzed by Orbitrap Fusion LUMOS Tribrid Mass Spectrometer, the precursors were ionized using an EASY-Spray ionization source (Thermo Fisher Scientific) source held at +2.0 kV compared to ground, and the inlet capillary temperature was held at 320 °C. Survey scans of peptide precursors were collected in the Orbitrap from 350-1800 Th with an AGC target of 400,000, a maximum injection time of 50 ms, RF lens at 30%, and a resolution of 120,000 at 200 m/z. Monoisotopic precursor selection was enabled for peptide isotopic distributions, precursors of z = 2-8 were selected for data-dependent MS/MS scans for 3 seconds of cycle time, and dynamic exclusion was set to 15 seconds with a ±10 ppm window set around the precursor monoisotope.

In EThcD scans, an automated scan range determination was enabled. An isolation window of 2 Th was used to select precursor ions with the quadrupole. An SA collision energy of 35%, an AGC target value of 50,000, a maximum injection time of 54 ms, and a resolution of 30,000 at 200 m/z were selected.

##### Data Analysis

Data processing was carried out using Thermo Proteome Discoverer 2.4 using a SwissProt homo sapiens database (TaxID=9606 and subtaxonomy, 42253 protein sequences). Carbamidomethyl (Cys) were chosen as static modification, oxidation (Met) and HexNAc (Ser or Thr) were chosen as variable modification. Mass tolerance was 10 ppm for precursor ions and 0.02 Da for fragment ions. Maximum missed cleavages was set as 2. Peptide spectral matches (PSM) were validated using the Percolator algorithm, based on q-values at a 1% FDR at both peptide and protein level.

#### Colony formation assay

H12299 and A549 cells stably expressing HA-YTHDF2-WT or -T49A plasmids were seeded in 10D plates (800 cells in each plate). After being cultured for 14 days, the cells were stained by Crystal violet. The plates were photographed and the cells were counted. All experiments were repeated three times and quantified.

#### Cell proliferation assay

H12299 and A549 cells stably expressing HA-YTHDF2-WT or -T49A plasmids were cultured in six-well plates for proliferation assay. Cell numbers was measured every 24 h and continuously measured for 72 h. All experiments were repeated three times and quantified.

#### Wound healing assay

H12299 and A549 cells stably expressing HA-YTHDF2-WT or -T49A plasmids were cultured in a six-well plate and then the cells were scratched with a 200 µl pipette tip when the cells almost 100% cover the well. Wound areas were photographed at 0 h and 48 h using a microscope. All experiments were repeated three times and quantified by ImageJ. Cellular migration rates were quantified by calculating the percentage of area reduction.

#### Transwell assay

H12299 and A549 cells stably expressing HA-YTHDF2-WT or -T49A plasmids were cultured in six-well plates and starved for 24 hours by using serum-free DMEM. 3 × 10^3^ cells were seeded into Transwell upper chamber (8-μm pore size; Millipore) with 50 μL serum-free DMEM, and the lower chamber was added with 200 μL DMEM with 10% FBS. Cells on the filter side of the chamber was stained by 0.1% Crystal violet at room temperature for 10-20 min after 24 h. All experiments were repeated three times.

#### Mouse xenograft analysis

For xenograft assays, 2 × 10^5^ HA-YTHDF2-WT or -T49A cells were resuspended in Matrigel (Corning) and then injected into the right abdomen of nude mice. The tumor volumes were measured every 3 days and being continuously measured from 7^th^ day to 25^th^ day. The mice were obtained from the Beijing SPF Biotechnology Co, Ltd (Certification NO. SCXK (Jing) 2019-0010). All animal work procedures were approved by the Animal Care Committee of Capital Normal University (Beijing, China).

## Abbreviations

ETD: Electron transfer dissociation
MS: Mass spectrometry
O-GlcNAc: O-linked β-N-acetylglucosamine
OGT: O-GlcNAc transferase
PTM: post-translational modification
IP: Immunoprecipitation
IB: Immunoblotting
TMG: Thiamet-G;
OGA: O-GlcNAcase
CHX: cycloheximide
YTHDF2: YTH domain family 2
ERK: Extracellular-signal regulated kinase
Ub: ubiquitin

## Acknowledgements

Jing L. is supported by the National Natural Science Foundation of China (NSFC) fund (32271285 and 31872720) and R & D Program of Beijing Municipal Education Commission (KZ202210028043). Y. G. is supported Non-profit Central Research Institute Fund of Chinese Academy of Medical Sciences (2021-RC350-002), CAMS Innovation Fund for Medical Sciences (2021-I2M-1-026).

## Data Availability

The mass spectrometry proteomics data have been deposited to the ProteomeXchange Consortium via the PRIDE (36) partner repository with the dataset identifier PXD045137 and 10.6019/PXD045137.

